# UVfinder: a tool to extract bryophyte sex-linked gene copies from the GoFlag408 probe set

**DOI:** 10.64898/2026.07.01.735932

**Authors:** Seyeon Kim, John L. Bowman, Edward L. Braun, Stuart F. McDaniel

## Abstract

**Premise:** Target enrichment sequencing using probe sets like GoFlag 408 has revolutionized phylogenetics, yet recent genomic data indicate that some probes may be sex-linked, potentially introducing topological conflict but also allowing studies of sex-specific evolutionary processes. To test for sex-linkage across the bryophytes, we developed UVfinder, a pipeline designed to identify sex-linked GoFlag loci across published moss genomes and enable sex-aware downstream analyses.

**Methods and Results:** Applying UVfinder to 50 dioicous moss genomes,we identified 93 probes that exhibit sex-linkage in one or more lineages. Our results provide genomic evidence for neo-sex chromosome formation via autosome-sex chromosome fusion and gene translocation. By comparing species trees derived from sex-linked versus autosomal loci in Hypnales and Dicranidae, we demonstrate that sex-linked loci harbor phylogenetic information that is distinct from that in autosomes. We also discovered a pervasive female sampling bias perhaps reflecting a preference among collectors for plants with sporophytes.

**Conclusions:** Our findings highlight the dynamism in sex linkage across bryophytes and suggest that sex-aware phylogenomics can be used to reconstruct ancestral karyotypes and potentially resolve topological conflict. We also expect that UVfinder will facilitate study of sex specific evolutionary processes, particularly with improved genome assemblies and increased sampling in males.

## INTRODUCTION

Target enrichment sequencing approaches, like the GoFlag408 probe set (Breinholt et al., 2021) have provided a cost-effective means to conduct phylogenomic analyses on a wide variety of sample types, allowing researchers to generate rich datasets from potentially degraded herbarium samples (Aguado□Ramsay et al., 2024; Bechteler et al., 2023; Budke et al., 2023; Draper et al., 2022; Jauregui-Lazo et al., 2023; Liu et al., 2019; Medina et al., 2019; Peñaloza-Bojacá et al., 2025). However, the use of multi-locus analyses has also forced researchers to confront the fact that different loci may support alternative phylogenetic relationships (Bechteler et al., 2023). In sexually reproducing organisms, patterns of variation in the genome represent not a single history but rather a multitude of potentially distinct histories. The stochastic segregation of ancestral variation into descendant species (i.e., incomplete lineage sorting), hybridization among species, or gene duplication and loss often cause genealogical conflict at multiple taxonomic levels, with consequences for studies of phenotypic evolution (Hibbins et al., 2023; Pease et al., 2016). Identifying the evolutionary processes that generate biodiversity requires explicitly accounting for the causes of topological conflict.

Among the bryophytes, including mosses, liverworts, and hornworts, topological conflict caused by sex-linkage may be both common and potentially easy to circumvent. About half of bryophyte species are dioicous (have separate males and females) and may have ancient sex chromosomes (Laenen et al., 2016; McDaniel et al., 2013; Villarreal & Renner, 2013). Bryophytes have a UV sex chromosome system in which both sexes have a sex-specific, non-recombining chromosome (the U is transmitted only through eggs, the V is transmitted only through sperm). The bryophyte U and V chromosomes each contain up to hundreds or even thousands of genes (Carey et al., 2021; Silva et al., 2021; Yu et al., 2022). Although the GoFlag408 set targets single-copy nuclear loci, it inadvertently included some genes with U and V-linked copies, which loci are sex-linked varies among lineages because the sex chromosomes are evolutionarily dynamic, occasionally fusing with autosomes (Carey et al., 2021; Yue et al., 2026). Analyzing diploid sporophyte DNA with highly diverged female and male orthologs can introduce phylogenetic error if target-capture probes match sex-linked genes. Reversions to monoicy are also very common and may result in the stochastic or biased loss of one or the other sex-linked copy (Singh et al., 2023).

The higher-level phylogeny of the bryophytes has remained relatively stable in recent years, although recent analyses of target-capture data have underscored the abundant and unexplained topological conflict at many nodes in the phylogeny (Bechteler et al., 2023; Liu et al., 2019). Given the observed support for conflicting topologies in the bryophyte tree (Bechteler et al., 2023), it is naïve to assume that simply acquiring more data is likely to resolve many critical nodes. However, theory suggests that sequence data from some genomic regions – for example, data from non-recombining sex chromosomes – will exhibit higher congruence with inferred species trees than recombining autosomal genes (Li et al., 2019; Pease & Hahn, 2013; Thom et al., 2024). Thus, while analyses of sex-linked target capture data from mixed-sex samples or sporophyte DNA are very likely to generate positively misleading genealogical information, in marked contrast, sex-aware analyses (i.e., those using known male and female plants, for dioicous species) of haploid sex-specific data are predicted to produce higher quality phylogenies. Analyses of known-sex haplotypes provide an independent means to reconstruct the history of sexual system (e.g., switches between dioicy and monoicy), sex chromosome-autosome fusions, and gene loss following transitions to monoicy, major questions in both the bryophytes and other groups. Additionally, targeting multiple female- and male-linked non-recombining genes provides the necessary genetic material to study sex-specific evolutionary processes, including mutation rates, demography, sex ratios, paternity, and population structure. In this paper, we describe a tool aimed at identifying sex-linked loci and generating preliminary phylogenetic analyses to compare the signal in sex chromosomes and autosomes. We focus on mosses, where more GoFlag loci are sex-linked, but we also provide a preliminary demonstration of the utility of the pipeline in liverworts and hornworts.

## METHODS AND RESULTS

To facilitate identifying bryophyte sex-linked genes in the GoFlag408 probe set, we developed a pipeline (UVfinder) that identifies taxon-specific sex-linked genes and generates a preliminary evolutionary analysis of the sex-linked genes. We have implemented UVfinder as a reproducible Snakemake workflow (Köster & Rahmann, 2012). Users provide genome assembly files as input. The pipeline then performs automated tBLASTn searches using sex-specific markers and the “GoFlag query set” as queries. This approach will allow inference of the sex of the individual used for each genome assembly and identify putative sex-linked loci. Next, the tool returns a list of sex-linked loci and finally constructs gene trees for the sex-linked loci. With these gene trees, users can identify gene duplications, which may arise for a variety of reasons and may or may not involve sex chromosomes-autosome fusions, filter out confounding sex-linked signals to reconstruct accurate species trees, and ultimately trace the evolutionary history of sexual systems, which can all be facilitated by the post-processing scripts provided in UVfinder.

### Usage

Figure 1 illustrates the overall workflow of the UVfinder pipeline. The UVfinder pipeline requires chromosome-level genome assemblies and is managed through two main configuration files. The *config/samples.tsv* file (Table S1) defines the metadata and paths for the input genomes, while the *config/config.yaml* file enables customizable parameter settings for bioinformatic tools. Then the pipeline renames the headers of all the genome files (Figure 1 Step 1) to facilitate accurate curation of genome files. With the renamed genomes, UVfinder constructs a BLAST database for each genome for later sex-markers, and GoFlag query set BLAST searches (Figure 1 Step2). Then, depending on the computer resources, UVfinder performs a BLAST search (McGinnis & Madden, 2004) using male/female sex markers (Figure 1 Step 3) and GoFlag Probe sets (Figure 1 Step 4). Male and female sex markers were queried against each genome using tBLASTn, and sex chromosomes were identified based on BLAST e-values and bit-scores. Hits with e-values > 1e−11 or bit-scores less than 85% of the maximum were excluded. For each genome, the chromosome was assigned as U (female-linked) or V (male-linked) according to which marker produced the higher bit-score (Figure 1 Step 5). Some genomes in Hypnales, liverwort, and hornwort have male and female sex marker BLAST hits on the same chromosome. Therefore, UVfinder performs well in identifying sex chromosomes, but sex of a genome could be difficult to determine definitively based solely on the highest bit-score rule.

**Figure 1.**
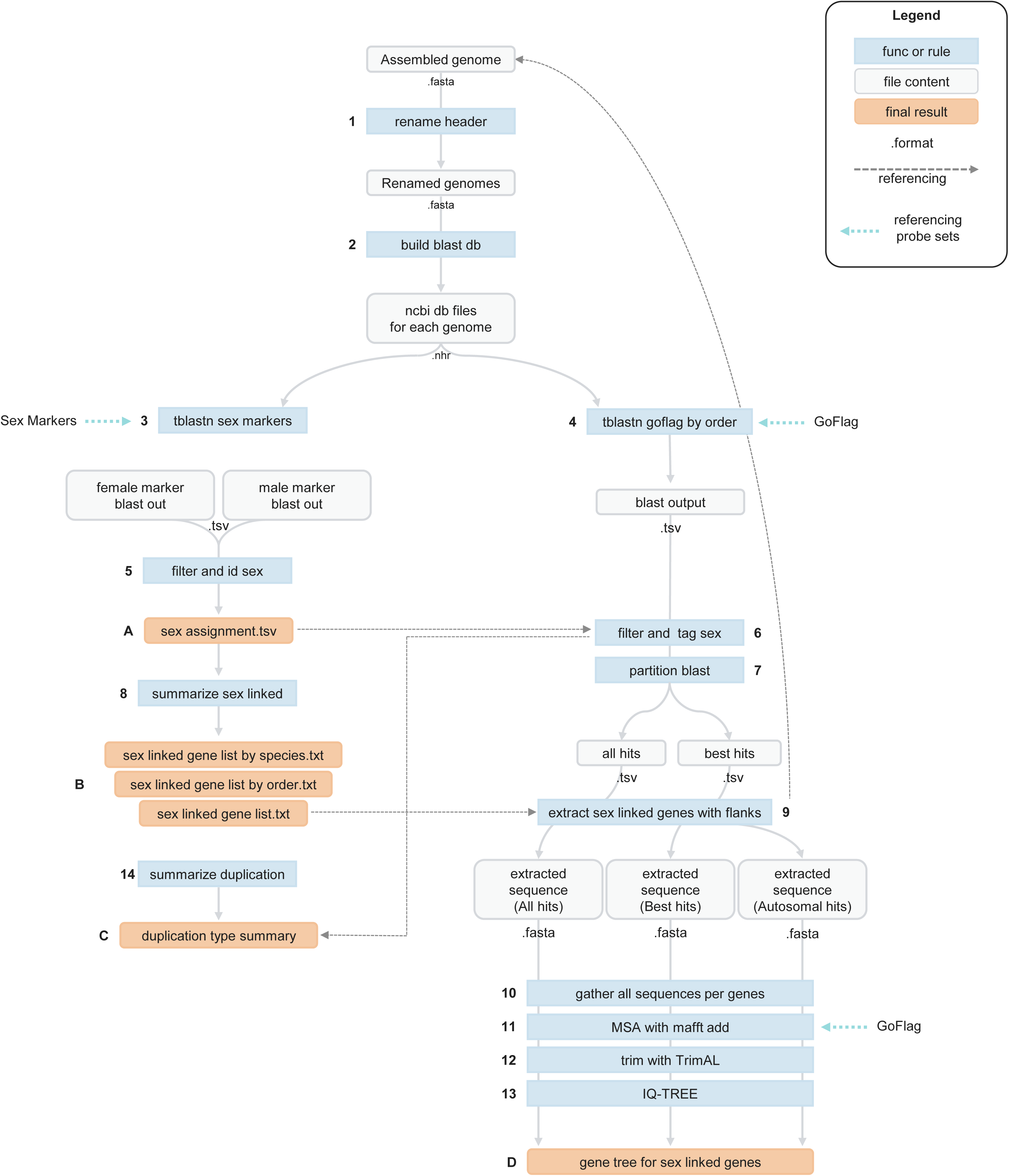
Overall workflow of UVfinder. A schematic flowchart illustrating the step-by-step pipeline. Light blue boxes represent processing functions or rules, gray rounded boxes indicate intermediate file contents with their corresponding file extensions (e.g., .fasta, .tsv, .nhr), and orange boxes denote the final key outputs. Numbers (1–14) indicate the sequential steps of the workflow, while letters (A–D) highlight major checkpoints and final results. Dashed arrows represent data referencing across different stages, and dotted teal arrows indicate the input of external probe sets (Sex Markers and GoFlag).

The pipeline performs tBLASTn searches using the translated (amino acid) sequences from the GoFlag query set against all input genomes. Because each GoFlag query comprises sequences from 531 moss taxa, the probe set was filtered by taxonomic order corresponding to each input species to improve alignment accuracy and reduce computational burden. When no matching order was available, the entire probe set was used without filtering. BLAST hits with e-values > 1e−15, probe identity < 70%, or bit-scores < 85% of the maximum were excluded. Filtered hits were labeled as “U” or “V” if they mapped to the sex chromosome, “A” if they mapped to autosomes, and “N” if the sex chromosome could not be determined using sex-specific markers (Figure 1 Step 6). After the GoFlag BLAST searches, UVfinder partitions the BLAST results into “Best_hits” and “All_hits” (Figure 1 Step 7). The “Best_hits” dataset retains a single GoFlag BLAST hit per gene, whereas “All_hits” includes all non-overlapping hits passing the filtering criteria to account for duplication events such as duplications in a single gene, or small chromosomal region, sex-linked duplication, and whole genome duplication. Sequences from “Best_hits”, “All_hits”, and “Autosoaml_hits (Not sex-linked hits in any of the input species)” were then extracted with a default flanking window of ±100000 bp (Figure 1 Step 9), and sequences were grouped by gene (Figure 1 Step 10). Then, sequence alignments were performed using MAFFT v7.526 with the GoFlag nucleotide dataset as a reference(--genafpair --maxiterate 1000 --adjustdirection) (Figure 1 Step 11) (Katoh & Frith, 2012). Alignments were trimmed with trimAl v1.5.1 using the parameters -automated1 to reduce noise (Figure 1 Step 12) (Capella-Gutiérrez et al., 2009). Phylogenetic trees for each sex-linked gene were reconstructed using IQ-TREE v3.0.1 with automatic model selection (“-m MFP”) and 1,000 ultrafast bootstrap replicates to infer sex-specific evolutionary patterns (Figure 1 Step 13) (Wong et al., 2026). Users can change this option through the configuration file. While we utilized these specific versions for our analyses, the pipeline is compatible with other versions of these programs.

In addition, based on the filtered BLAST output, duplicated gene pairs were categorized into specific duplication types according to their genomic location and sex chromosome assignments (Figure 1 Step 14; Figure 3A). Duplications were first assessed by location (on the same chromosome or across different chromosomes). For intra-chromosomal pairs, a user-adjustable distance threshold of 100,000bp was applied to separate local duplications from broader intra-chromosomal duplications. Inter-chromosomal events were divided into three categories depending on the nature of the chromosomes involved (between two autosomes: Inter-chromosomal Duplication, between an autosome and a sex chromosome: Sex-linked Duplication, or between two uncharacterized chromosomes: Inter-chromosomal Duplication in Sex Unknown Samples)

### Data acquisition

We used the comprehensive phylogenomic dataset from Bechteler et al. (2023), which we call as “GoFlag query set”. This dataset was generated using the GoFlag 408 flagellate land plant probe set, which targets 405 exons representing 228 single- or low-copy nuclear genes. We retrieved the pre-aligned nucleotide and amino acid sequences for 531 bryophyte species from the associated Dryad Digital Repository (https://doi.org/10.5061/dryad.3j9kd51qm). The amino acid sequences were used as queries for homology searches with tBLASTn, while the aligned nucleotide sequences served as templates for subsequent multiple sequence alignments using “*mafft -- add*” (Katoh & Frith, 2012).

For the male sex marker, we use the V-linked copy of the Zinc Finger Ran binding protein from *Ceratodon purpureus*, representing the most ancient coalescence between U and V chromosomes reported by Cary et al. (2021) and exhibiting sex-linkage across all species descending from the common ancestor of *Buxbaumia* and *Ceratodon.* For the female sex marker, we initially considered the U-copy of the same Zinc Finger Ran binding protein. However, ultimately, we selected the *Marchantia polymorpha RKD* gene (Mp*RKD*), which is an *RWP-RK* transcription factor essential for egg cell identity (Bowman & McDaniel, 2026; Rövekamp et al., 2016), as it consistently maps to chromosomes with sex-chromosome characteristics (low gene density, high trnasposable element content) but not to chromosomes with male markers and yields more consistent results in identifying sex chromosomes and sex of the species.

We compared the performance of Mp*RKD* and U-copy of the Zinc Finger Ran binding protein as female sex marker and ultimately selected Mp*RKD*, *since* it consistently aligned better with findings from previous studies. Unlike the U-copy, which was prone to false positives or failed to assign sex chromosome, Mp*RKD* correctly identified U chromosomes in *Polytrichastrum alpinum* (chromosome 8; Zeng et al., 2025) and *Syntrichia caninervis* (chromosome 13; Silva et al., 2021). Furthermore, because *RKD* is exclusive to female or hermaphroditic individuals, its absence reliably indicates a male genome (e.g., *Sphagnum* except *S. balticum* and *S. palustre*, Figure S1).

### Use case example

To evaluate the utility of UVfinder, we used 50 dioicous moss genomes representing 49 moss species available on the NCBI genome repository (Table S1, accessed in Nov 2025), encompassing a diverse range of taxa and including genomes from both male and female plants. We found a total of 93 instances of sex-linkage among the GoFlag genes, with a range of 0 – 29 genes in a single genome. We first examine explanations for the variation in the counts of sex-linked genes among moss lineages. Next, we examine the patterns of shared sex-linked genes among lineages. We use the genomic location of newly sex-linked genes in related lineages where they are not sex linked to reconstruct karyotypic changes involving sex chromosomes. Finally, we compare the phylogenetic information in sex-linked and autosomal genes in groups where four or more taxa share a sex-linked gene. Throughout, we highlight both biological processes and data quality issues that could generate the observed patterns, provide use cases for the existing data, and new kinds of data that could improve the quality of the inferences regarding sex-linkage in mosses.

#### Sex-linked loci in the GoFlag query set across mosses

We found the most sex-linked genes in the Hypnales, Polytrichales, and Ditrichales, and the fewest in the Rhizogoniales, Orthotrichales, Bartramiales, and Sphagnales. Not surprisingly, we found a positive correlation between the number of samples analyzed and the total number of sex-linked genes identified, indicating that the number of sex-linked genes is strongly influenced by the number of genomes in the dataset (Figure 2B). This relationship is driven by the Hypnales, which had the highest number of both genomes (21 genomes) and sex-linked genes (55 genes). However, the orders Ditrichales (represented by a male/female pair of *Ceratodon purpureus*) and Polytrichales have a comparatively large number of sex-linked genes, even though we surveyed only a few genomes. The number of sex-linked genes we detected could reflect multiple processes, starting with variation in the sex chromosome gene content across species, either due to chance or alternatively could result from sex chromosome size variation among lineages. For example, *C. purpureus* possesses physically large sex chromosomes (∼110 Mbps), possibly because it experienced recent sex chromosome – autosome fusions (Carey et al. 2021). In contrast, the low gene count in the Sphagnales is consistent with the small sex chromosomes in the genus (∼5 Mbps) (Healey et al., 2023).

**Figure 2.**
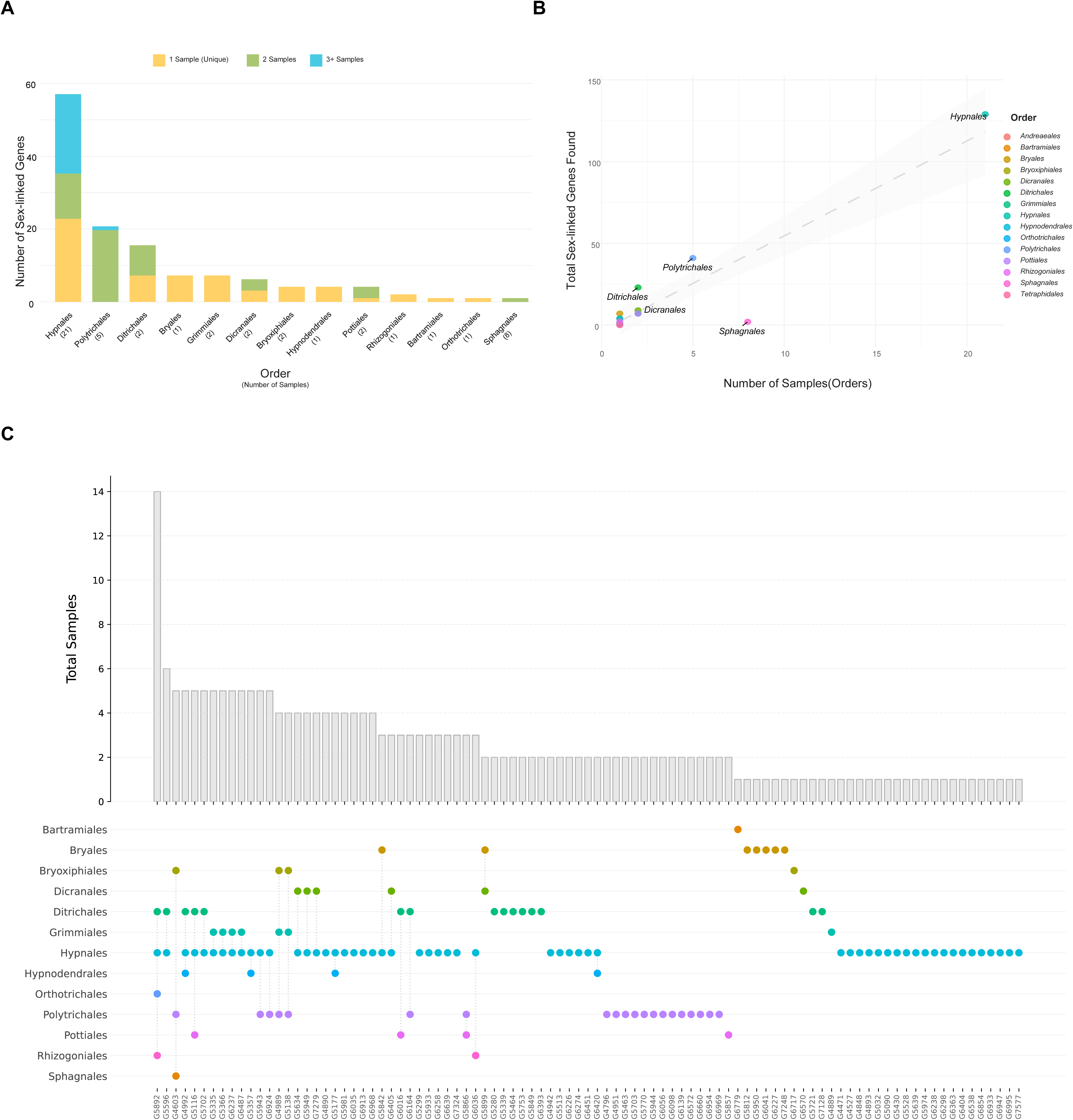
Sex-linked genes in 51 dioicous moss genomes. (A) Sample size for each order and the number of sex-linked genes in specific samples; (B) Distribution of sex-linked genes across moss orders; (C) Sex-linked gene counts in 51 dioicous moss samples and their distribution within 13 moss orders

It remains possible that the absence of sex-linked genes in some orders, such as Andreaeales, and Tetraphidales, reflects technical limitations with the input data. If the assembly is poor or fragmented, the pipeline may assign a scaffold or contig as a sex chromosome, meaning the pipeline would not detect linkage to GoFlag loci on the chromosome (this may be the case in the Tetraphidales; Mp*RKD* mapped to a scaffold). The pipeline also currently does not work equally well for all mosses because the *Ceratodon* Zinc Finger Ran binding protein, which we use as a male marker, was not sex linked in the common ancestor of all mosses; the U-V divergence of this gene occurred after the divergence of Sphagnales and the remaining mosses. Therefore, the Zinc Finger Ran binding protein cannot be reliably mapped to male Sphagnales sex chromosomes. We suspect that Sphagnales samples lacking the female marker (Mp*RKD*) are likely to be male, but without a positive male marker match, the current sex chromosome assignment algorithm categorizes them as ‘Unknown’. Therefore, beyond their small size alone, the low number of sex-linked genes observed in Sphagnales may be distorted by uncertainty in identifying males. Using the pipeline, Mp*RKD* only mapped to a chromosome in *Sphagnum balticum* and *Sphagnum palustre*, which have one common sex-linked gene, *G4603.* Based on the study of Sphagnales sex chromosomes (Healey et al., 2023), we found that the sex-linked gene (*G4603*) shared by *S. balticum* and *S. palustre* is also present on the sex chromosomes of other Sphagnales samples (Figure S1). To investigate its potential function, we BLASTed *G4603* against the reference genomes of both *Arabidopsis thaliana* and *Physcomitrium patens*. Interestingly, the sex-linked locus showed significant sequence homology to the THO complex subunit 5A (THO5A) in both species. THO complex consists <u>tr</u>anscription and <u>ex</u>port (TREX) complex (Ettner-Sitter et al., 2026), which together regulate female germline specification by repressing it in the plant ovule, ensuring that only a single megaspore mother cell (MMC) is normally formed (Su et al., 2017). While the functional annotation of *G4603* is suggestive, linking the gene to sexual function in other Sphagnum species requires analysis beyond the scope of this paper.

#### Shared sex-linked loci across species

Most of the sex-linked genes we found were unique to a single sample (Figure 2A), hough 17.3% were found in 3 or more genomes within the same order (Hypnales). Unlike other orders where pairs of shared genes were genus-restricted, (Pottiales and Polytrichales), the shared sex-linked genes in Hypnales are observed across diverse genera. This may be because different generic names are used at shallower genetic divergences in the Hypnales than in other bryophyte orders or recurrent sex chromosome – autosome fusions.

Data from the Polytrichales provide evidence for the recruitment of new sex-linked genes, most likely following a neo sex chromosome formation. In Polytrichales, 20 sex-linked genes were shared between *Polytrichum commune* and *Polytrichum formosum*, but they are autosomal in other dioicous Polytrichales species. Because the newly sex-linked loci are together on a chromosome in a sister species, it is likely that the loci were not translocated one by one, independently, but rather represent the translocation of a whole chromosome, although, without better outgroup information, we cannot exclude the possibility that the condition in the sister group represents a sex chromosome fission.

The gene that was sex-linked in the most orders was *G5982* (protein of unknown function), which appears to have independently translocated to the sex chromosome in the Ditrichales, Hypnales, Orthotrichales, and Rhizogoniales (Figure 2C). While Sphagnales and Polytrichales exclusively retain autosomal copies (A), we found sex-linked copies (U/V) in some Hypnales species, including male and female genomes, *Aulacomnium turgidum* (Rhizoginales), and *Orthothecium rufescens* (Orthotrichales), and the gene tree has high topological conflict (Figure S1). In *C. purpureus*, there were multiple inter-autosomal duplication events both in male and female genomes, and a sex-linked duplication in the female genome. The dispersed pattern of sex linkage likely reflects multiple independent events of parallel recruitment into the UV sex chromosomes via inter-chromosomal fusions or gene transpositions (Figure S1). Collectively, these observations highlight the complex karyotypic variation that can occur even in genes that were selected for the GoFlag probe set for their sequence conservation and low copy number.

#### Genome-wide gene duplications across mosses

To estimate the frequency of duplication in sex-linked genes and autosomal genes, we classified duplicated loci into distinct categories based on their chromosomal context: local (<100.000 bp), intra-chromosomal (>100,000 bp), inter-autosomal, and sex-linked duplication (Figure 3A).

**Figure 3.**
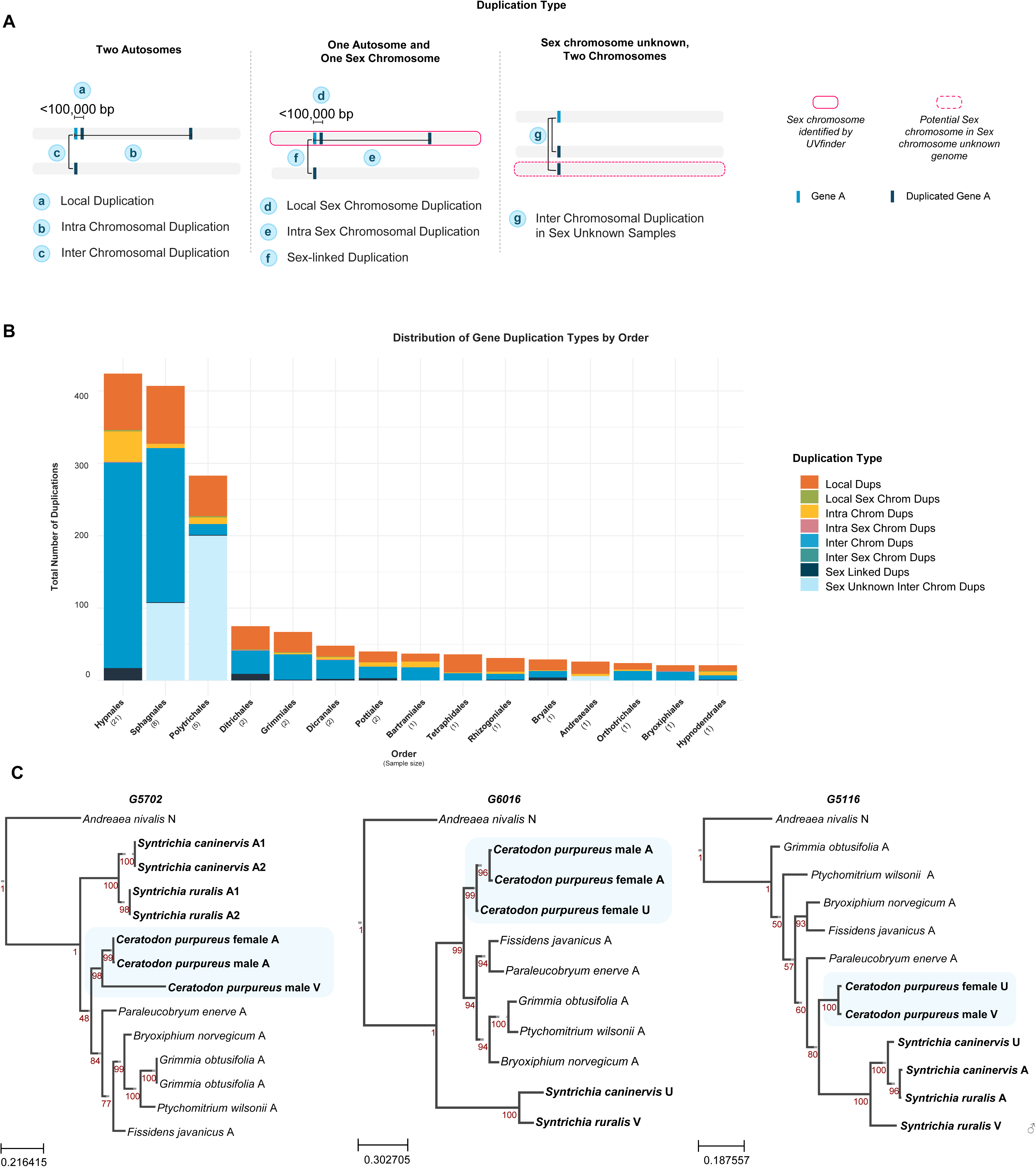
Duplication analysis on 51 dioicous moss genomes. (A) Duplication type based on genomic region; (B) Distribution of gene duplication types by order; (C) sex-linked duplication examples.

Across the 50 moss genomes, sex-linked duplications are rare (Figure 3B). Most duplicated genes are autosomal, though sex-linked duplications are present in Ditrichales, Pottiales, and Bryales, despite their small sample size. Conversely, in orders like Sphagnales and Polytrichales, many duplication events fall into the “Sex Unknown Inter Chrom Dups” (Figure 3B, light blue bars). This is likely driven by putative polyploidy and the lack of male markers in these lineages; that is, many sex-specific scaffolds remain unassigned, potentially masking true sex-linked duplications.

Importantly, in our input dataset, only *Ceratodon purpureus* has both a male and a female genome, which allows insights into the sex-specificity of duplication events. At locus *G5702*, the female genome retains a single autosomal copy, whereas the male genome harbors two copies, where one is on the autosome and the other is on the sex chromosome. The autosomal copies are alleles of the same locus, sister to the male-specific **V**-linked copy, suggesting a recent duplication subsequently translocated either onto or off the **V** chromosome. Because *Grimmia* and *Syntrichia* also have autosomal copies of *G5702*, interestingly also duplicated, we suspect that the translocation in *Ceratodon* was likely recent and onto the V chromosome. The long branch of V-copy suggests rapid sequence divergence following the duplication. Conversely, for locus *G6016*, the male genome has an autosomal copy, while the female genome possesses two, one U-linked and one autosomal. The short branch, compared to the V copy of *Ceratodon G5702*, suggests that there was a recent duplication or gene translocation onto U chromosome. In *Syntrichia caninervis and Syntrichia ruralis, G6016* is on both the U and V, respectively. This shows that the ancestral state of the *G6016* was autosomal, and it was independently recruited to the sex chromosome via a female-specific duplication in *Ceratodon* and potential chromosome fusion in *Syntrichia* U/V sex chromosome. In contrast to loci *G5702* and *G6016,* locus *G5116* lacks an autosomal paralog in *Ceratodon,* instead harboring sex-linked copies on both the U and V. In *Syntrichia*, both autosomal and sex-linked copies are present. Notably, the autosomal copies of *S. caninervis* and *S. ruralis* cluster together as sister taxa, while their available sex-linked copies (U in *S. caninervis* and V in *S. ruralis*) coalesce earlier. This topology, where orthologous autosomal copies group closer to each other than to their respective sex-linked paralogs, suggests that the duplication and subsequent recruitment to the sex chromosomes occurred in the common ancestor of these *Syntrichia* species, before their divergence.

#### Reconstructing karyotypic evolution in the Hypnales

To illustrate some of the challenges of identifying sex-linked genes in mosses using the GoFlag probe set, and to provide examples of how to avoid common pitfalls, we work through a reconstruction of the karyotypic evolution in the comparatively well-sampled Hypnales genomes, focusing on the sex chromosomes (Figure 4). We mapped the GoFlag BLAST results onto the assembled chromosome and colored the shared sex-linked genes in Hypnales. Based on this phylogeny and the chromosome map, we infer that the common ancestor of Hypnales clade possessed a karyotype of n = 10 + U/V, with the sex chromosomes containing gene *G5892*. Notably, our analyses tell us that *G5892* is not critical for sex-specific function but may be linked to loci that can easily evolve sex-specific functionality. In multiple Hypnales genomes, *G5892* is translocated to other autosomes (Figure 4, event b). For example, in *Heterocladium heteropterum*, the ancestral karyotype is conserved, but *G5892* is translocated to an autosome without whole-chromosome fusion or fission. In this case, the original sex chromosome maintains its identity as the sex chromosome. Using this ancestral state as a baseline, we identified five distinct structural events driving sex chromosome turnover.

**Figure 4.**
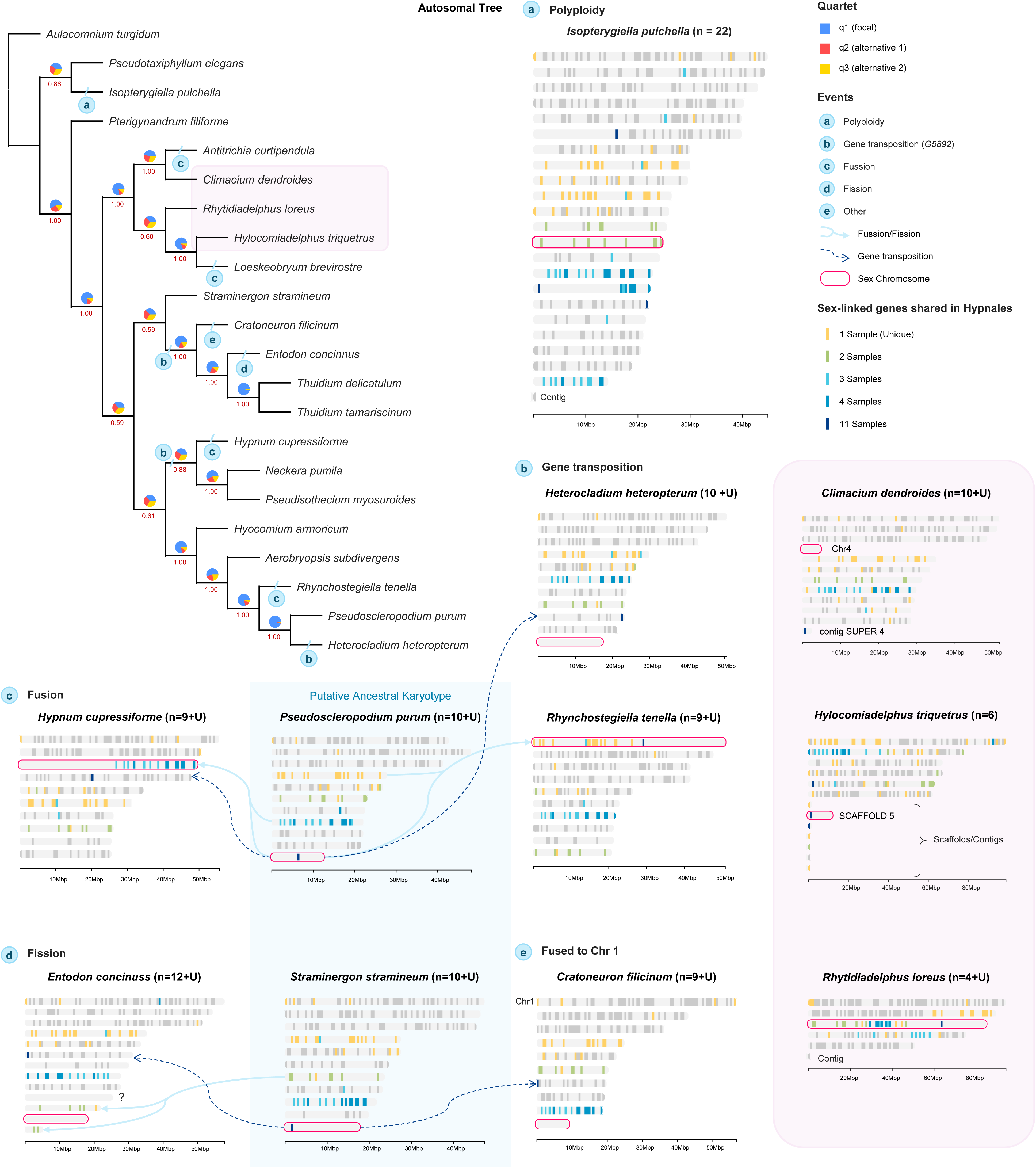
Evolutionary dynamics of karyotypic changes in Hypnales. Structural variations mapped onto the maximum likelihood autosomal gene phylogeny. Numbers below branches indicate bootstrap support, and multicolored pie charts represent quartet support (q1: focal topology, q2: alternative 1, q3: alternative 2) to evaluate phylogenetic conflicts. Circled letters (a–e) pinpoint specific genomic events along the branches, corresponding to chromosome map examples panels. Gray horizontal bars represent individual scaffolds or chromosomes scaled in megabases (Mbp). Gray colored bars on chromosome map denote GoFlag probe hits, with colors indicating the number of sharing samples (ranging from 1 to 11). Pink solid outlines represent sex chromosomes identified by UVfinder.(a) Polyplidy: recent polyploidy event in Isopterygiella pulchella (n = 22). (b) Gene transposition: Inter-chromosomal transposition of a specific sex-linked locus (G5892). (c) Fusion: Chromosomal fusion events leading to altered karyotypes. Light blue arrows indicate the directional fusion of chromosomal fragments.(d) Fission: Chromosomal fission events observed in *Entodon concinnus* (n=12+U), with light blue arrows showing fragment division. Fission cannot solely explain the karyotypic changes in *E.concinnus*, where we marked with the question mark. (e) Other events: In *Cratoneuron filicinum*, chromosome fusion happened after the translocation of G5892. However, we treated this case separately because we could not identify the chromosome fused to the chromosome 1. Genomes that have an unusual pattern of sex chromosome *(Climacium dendroides* (n=10+U)) or possess much smaller number of chromosomes (*Hylocomium triquetrus* (n=6) and *Rhytidiadelphus loreus* (n=5)) are indicated with pink box both on the tree and the chromosome map.

First, we identified a recent whole-genome duplication (WGD) event in *Isopterygiella pulchella* (n = 22; Figure 4, event a). This event is supported by 195 inter-chromosomal duplication events among GoFlag query loci, including nine sex-linked duplications. Mapping these inter-chromosomal duplications onto the *I. pulchella* chromosomes using the Circos R package revealed syntenic blocks (Figure S3). This recent WGD event is further corroborated by paralogous gene pairs splitting at the terminal tips of individual gene trees. In bryophytes, polyploidy is frequently coupled with transitions from dioicy to monoicy. However, the sexual system of *I. pulchella* remains uncertain. While the Flora of North America (2014) and British Bryological Society (2014) characterize it as autoicous (monoicous), Ignatova et al. (2020) suggested it might be autoicous or potentially dioicous with yet-undiscovered male plants. Our data reflects this ambiguity. The female-specific sex marker Mp*RKD* mapped to two locations, chromosome 13 and chromosome16, potentially reflecting a recent duplication, while the male-associated *Ceratodon* Zinc-finger Ran binding protein mapped to chromosome 6. The retention of both U and V chromosomes in a polyploid may be common in transitions to monoicy (Singh et al., 2023). Therefore, the species may be monoicous and possess only ancestral U and V chromosomes, with inheritance patterns that are not currently correlated with sex.

Interestingly, the *G5892* locus, which is often sex-linked within Hypnales, is duplicated across multiple genomic regions. Both chromosome 16 and chromosome 6, which based on our marker data are the candidate ancestral U and V, respectively, harbor duplicated copies of the sex-linked gene *G5892,* but so did chromosome 17, which lacks any sex markers. Interestingly, the highest-scoring BLAST hit for the female marker mapped to chromosome 13, where *G5892* was absent. This underscores the challenges posed by polyploidy when using the UVfinder, as homeologous or duplicated sex-linked regions and overlapping male and female markers can confound the sex chromosome identification step of the pipeline.

Next, we examine the reduction in chromosome count (n = 9 + U) in *Hypnum cupressiforme* and *Rhynchostegiella tenella* (Figure 4, event c). Our mapping demonstrates that each of these reductions resulted from the fusion of an autosome with the ancestral sex chromosome, thereby forming a neo-sex chromosome. While the exact mechanism is unknown, theory and phylogenetic studies suggest that such fusions may be adaptive under some circumstances (Anderson et al., 2020). This newly formed neo-sex chromosome can arise either from the fusion of an autosome with the ancestral common sex chromosome, as seen in *R. tenella*, or via an alternative autosomal fusion. In contrast to this chromosome loss, a gain in chromosome number (n = 12 + U) was observed in *Entodon concinnus* (Figure 4, event d). This indicates that a chromosomal fission event occurred, although this fission alone does not fully explain the observed numerical changes. The chromosome assembly quality was good enough, therefore, necessitating further comprehensive analysis. In *Cratoneuron filicinum* (n = 9 + U), *G5892* translocated to the autosome prior to the node where *C. filicinum* and its sister group diverged (Figure 4, event e). In addition, a chromosome reduction event occurred via the fusion of an unidentified autosome with chromosome 1, which subsequently expanded the size of chromosome 1 by approximately 10 Mb. However, given the limited number of genes and coverage afforded by the GoFlag query set, we cannot identify the exact fusing partner chromosomes.

We report unusual genome assembly characteristics in *Climacium dendroides*, *Hylocomiadelphus triquetrus*, and *Rhytidiadelphus loreus*. Despite initial concerns regarding assembly quality, their assembly statistics were good. The reference genome of *C. dendroides* comprises 11 chromosome-level scaffolds, aligning with the standard Hypnales karyotype. However, chromosome 4 (initially named SUPER_4) exhibits an exceptionally reduced size compared to the autosomes and other Hypnales sex chromosomes. Bell and Long (2024) noted that chromosome 4 likely represents a U or V sex chromosome. The DNA used for assembly originated from different individuals of unknown sex. The absence of Hi-C signals on chromosome 4 indicated that this chromosome was not present in the individual used in this extraction. Our UVfinder analysis clarifies this ambiguity, indicating that the absence of the Hi-C signal reflects a sex mismatch. Specifically, the female-specific marker Mp*RKD* and the gene *G5892* both mapped to chromosome 4, providing empirical evidence that chromosome 4 is the U sex chromosome.

*H. triquetrus* and *R. loreus* possess n = 6 and n = 5 chromosomes, respectively, retaining most of the GoFlag query hits. The presence of most of the GoFlag loci suggests that the reduction in chromosome number resulted from chromosomal fusions (i.e., descending dysploidy). These observations show that BLAST-based mapping provides preliminary insights into karyotypic evolution, although a more rigorous, high-resolution synteny analysis with more complete taxonomic sampling is required to confirm such structural rearrangements. While the fusions, fissions, or translocations could accurately represent the karyotypic change, they may also reflect genome assembly artifacts. The GoFlag mapping approach provides a computationally simple way to identify homologous chromosomal segments in genomes that lack annotations, thereby identifying important nodes for more focused analyses.

#### Phylogenetic signal in sex-linked and autosomal loci

We originally designed the UVfinder to identify sex-linked markers for population genetic studies, and because sex-aware analyses are expected to produce phylogenies more consistent with the species tree than naive analyses of all GoFlag loci. To test this, we compared species trees generated from the complete GoFlag dataset (“All genes”), trees constructed from genes that are sex-linked at least once in the Hypnales (Figure 5A) and Dicranidae (Figure 5B) (“Sex-linked”), and trees constructed exclusively from autosomal genes in the input clades (“Autosomal”). For each dataset, we estimated individual maximum likelihood (ML) gene trees. To account for gene-tree estimation error, we collapsed unstable internal branches with bootstrap support values below 10% into polytomies (Zhang et al., 2018). Finally, we pooled these collapsed gene trees to infer the species tree using ASTRAL-pro3 (Zhang et al., 2020). Topologies of the “All genes” and “Autosomal” trees were highly congruent in both Hypnales and Dicranidae, indicating that sex-linked loci caused minimal topological conflict. Although the “Sex-linked” trees largely mirrored this topology, they exhibited different alternative topologies at some nodes.

**Figure 5.**
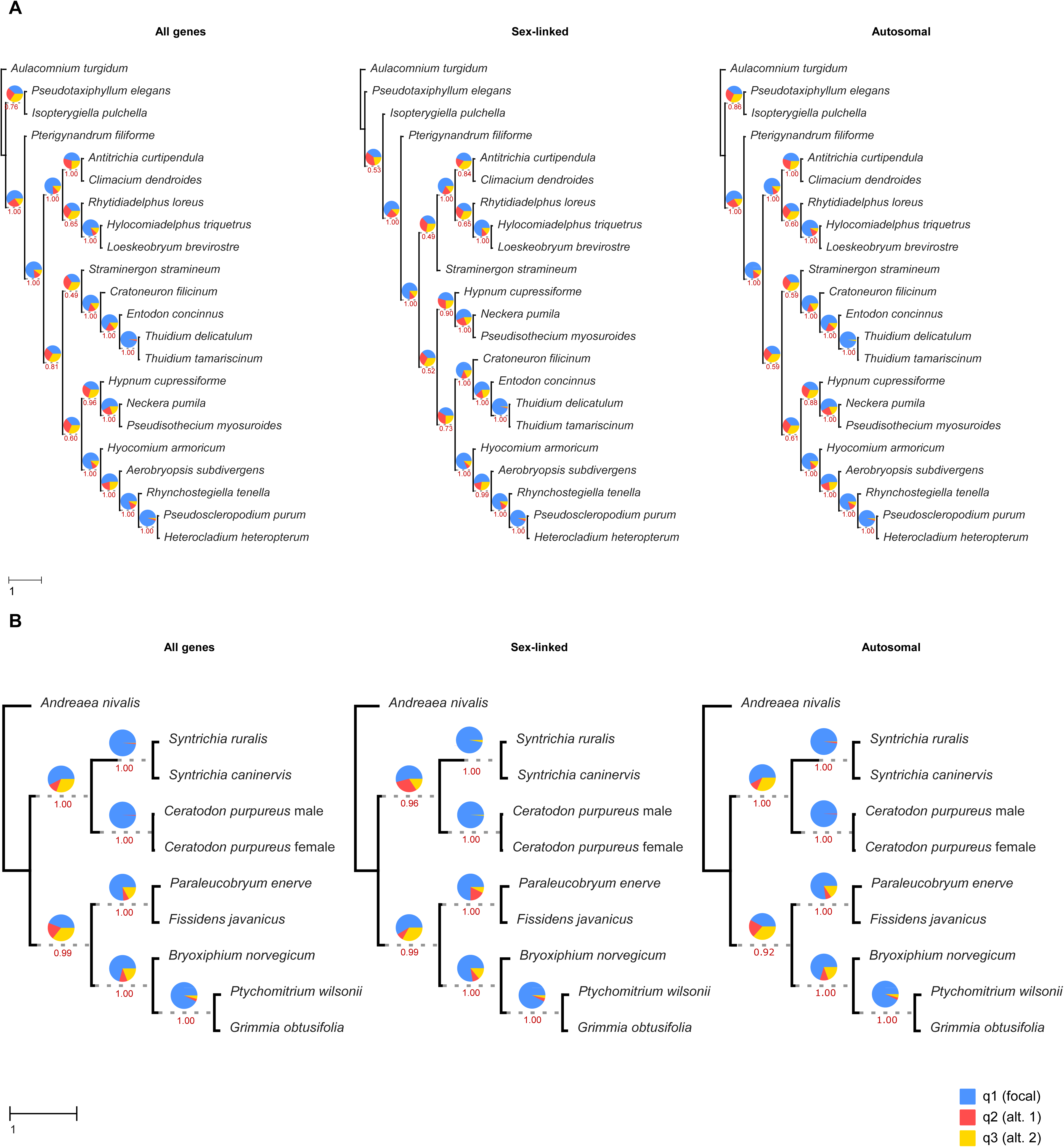
Comparison of phylogenetic discordance across different genomic datasets (All genes, Sex-linked genes, and Autosomal genes from GoFlag query set). Maximum likelihood and coalescent-based species trees reconstructed from three distinct datasets—All genes, Sex-linked genes, and Autosomal genes—to evaluate topological conflict. (A) Phylogenomic trees inferred for the order Hypnales and outgroup *Aulacomnium turgidum*. (B) Phylogenomic trees inferred for Dicranidae and outgroup Andreaea nivalis. Red values below branches indicate bootstrap support, while multicolored pie charts at each node represent quartet support scores based on ASTRAL-Pro3 analysis. Pie charts at the nodes represent the proportion of gene trees that support the main topology (blue), support the first alternative topology (red), support the second alternative topology (yellow). Scale bars represent branch lengths in coalescent units (CU).

These conflicting topologies suggest that sex-linked loci may have a distinct evolutionary history, though identifying the causal processes requires more data. Sex-linked loci are predicted to experience less ILS due to smaller effective population sizes and because they are more likely to host genes that contribute to post-zygotic reproductive isolation (Brandvain et al., 2014; Langdon et al., 2024; Li et al., 2019; Pease & Hahn, 2013; Schumer et al., 2018; Thom et al., 2024). Sex-linked loci are also subject to differential introgression (Fraïsse & Sachdeva, 2021). However, empirical evaluation of phylogenetic signal on sex chromosomes suggests a complex situation. For example, Rivas-González et al. (2026) found that the sex chromosome signal was generally more discordant than the autosomal signal, though details were method-sensitive. Finally, the complex history of the sexual system potentially confounds simple expectations.

In Hypnales and Dicrinidae, “All genes” dataset generally yields higher branch support values, with no drastic quartet score shifts at most nodes. This higher support likely reflects the larger number of genes used in the ASTER analysis (228 genes for “All gene trees and 173 genes for “Autosomal” tree in Hypnales, and 195 genes for “Autosomal” tree in Dicrinidae). Topological discordance from highly divergent sex-linked genes was minimal because our dataset is largely derived from the female genomes (20/21 genomes in Hypnales and 7/9 genomes in Dicrinidae), meaning few V-specific signals were present to cause phylogenetic conflict. Although branch support in some nodes in Hypnales was lower than in the “All genes” and “Autosomal” trees, which may be caused by the smaller number of genes used to construct the tree, the distinct topology of the sex-linked tree highlights that these loci harbor a unique evolutionary signal that deviates slightly from the autosomal signal. While accurately interpreting these patterns requires a detailed understanding of the historical transitions in sexual systems (e.g., between dioicy and monoicy), these conflicting sex-linked loci can provide valuable information for reconstructing the evolutionary history of sexual systems in these mosses

These findings demonstrate that the GoFlag genes are robust to the inclusion or exclusion of sex-linked signals, confirming their broad utility for phylogenomic studies across bryophytes. However, the topological shifts observed in the sex-linked trees underscore that incorporating sex-aware analyses remains crucial. Although filtering out sex-linked loci did not alter the major topology of the “Autosomal” species tree in these two cases, the topological shifts and alternative quartet frequency reversals show that ignoring sex linkage can still introduce topological conflict. The high congruence in both cases highlights how a female-biased dataset can mask sex-linked conflict by reducing V-specific signals. This female sampling bias may arise because researchers preferentially collect easily identifiable sporophyte-bearing (female) plants, inadvertently enriching for female DNA. Since phylogenetic signals depend heavily on the sex composition of input genomes, incorporating haploid, known-sex data is crucial for reconstructing high-quality phylogenies.

#### UVfinder application for liverworts and hornworts

While our primary focus is on mosses, UVfinder can successfully identify sex-linked GoFlag loci in liverworts and hornworts, demonstrating the broad utility of the tool. In liverworts, BPCU and BPCV are located on the U and V chromosomes, respectively, therefore, we used them as sex markers and identified sex-linked genes in eight liverwort species (Table S2). UVfinder successfully assigned sex chromosomes and sex-linked genes. However, the number of identified sex-linked genes in these species was low, ranging from 0 to 2 genes. This likely reflects their small physical size of the sex chromosomes in liverworts (approximately 5-10 Mbp), which naturally harbor fewer genes, as was the case for Sphagnales. In other cases, putative sex chromosomes were assigned to scaffolds or contigs rather than fully assembled chromosomes (e.g. *Lunularia cruciata, Conocephalum conicum,* and *Blasia pusilla*). As we noted above, highly fragmented assemblies can hinder the accurate identification of sex-linked genes. Furthermore, we observed discrepancies where sex chromosome assigned by UVfinder differs from the putative sex chromosome inferred by the chromosome size. For example, in *Riccia fluitans,* previous studies (Fu et al., 2025; Levins et al., 2025) reported that BPCU/V maps to an autosome, whereas *FGMYB* maps to chromosome 5, which is a putative sex chromosome. Our pipeline also mapped chromosome 2 as a sex chromosome using BPCU/V sex markers. Additionally, in *Ptilidium cilicare, Bazzania trilobata, and Plagiochila carrigtonii* the BPCU/V markers did not map to the smallest chromosomes, suggesting further investigation is needed. We should caution that BPCU and BPCV have high enough sequence similarity such that both markers may map to the same genome, complicating the assignment of an individual’s specific sex. While this ambiguity does not impede the identification of sex chromosomes and sex-linked genes, researchers conducting sex-specific studies must be cautious when determining sex based solely on these markers.

*FGMYB* is a key transcription factor that is a candidate female sex-determining gene in hornworts, (Bowman & Levins, 2025) and it is located on the ancestral female (U) sex chromosome. Therefore, we use hornwort *FGMYB* sequences as a female sex marker. However, we currently have no known ancient male sex-determining gene in hornworts. Therefore, we use V-linked *TCP1* and *C4HDZ* as male markers, which are conserved transcription factors that retain distinct gametologs on both sex chromosomes (Schafran et al., 2025). Since there are multiple female and male sex marker sequences from different species and genes, the pipeline identifies the most suitable male and female markers based on e-value and bit-score before comparing the male and female markers. We tested with three chromosomal-level assembled hornwort genomes publicly available (Table S3) and found no sex-linked genes in *L.dussii* and *P.chilonesis* and six sex-linked genes in *Phymatoceros. phymatodes*. In *Leiosporoceros dussii* and *Phaeomegaceros chiloensis,* the sex chromosomes inferred by UVfinder have a small size (7 and 9 Mbp each), with no GoFlag query set maps to the chromosome. However, in *P. phymatodes*, where the sex chromosome inferred by UVfinder has a small number of sex-linked GoFlag genes compared to its chromosome size. Interestingly, chromosome 5 is small (3Mbp), but still has 21 genes (Table S4). This disparity between the large physical size of the inferred sex chromosome in *P. phymatodes* and its low number of conserved GoFlag genes suggests a massive accumulation of transposable elements, which is a characteristic of non-recombining sex chromosome (Bowman & Levins, 2025; Carey et al., 2021; Yue et al., 2026). Furthermore, because GoFlag targets highly conserved loci, rapidly evolving or degenerated sex-linked genes may not be detected, resulting in an underestimation of the actual gene content. In addition, large number on the much smaller chromosome 5 (3Mbp, 21 genes) could indicate a gene-rich autosome, or it may be the true sex chromosome (or a gene-dense pseudo-autosomal region), and UVfinder misidentified another massive chromosome as the sex chromosome due to the incidental transposition of marker sequence. Therefore, higher-quality chromosomal assembly genomes are needed to identify and compare lineage-specific shared sex-linked genes.

## CONCLUSIONS

The GoFlag408 probe set was designed to support phylogenomic analyses in flagellate plants, but in combination with new genomic data, the tool can also be used to study sex-specific evolutionary processes in bryophytes. To facilitate such analyses, we developed UVfinder, an analytical pipeline designed to map the GoFlag408 target enrichment set against published moss genomes to identify and conduct preliminary molecular evolutionary analyses of sex-linked loci. We have shown how the pipeline can generate chromosome-scale visualization to investigate autosome-sex chromosome fusions and infer ancestral karyotypic states, detect and classify gene duplication events, specifically local and sex-linked duplications, and, by generating and parsing gene trees to identify the coalescence between U-V or other gene copies, provide a robust framework to pinpoint the precise evolutionary timing of major karyotypic changes and reconstruct the ancestral state. We provide worked examples of how UVfinder allows users to filter out sex-linked paralogs to construct phylogenies with less topological conflict that may be more consistent with the species tree. We expect that the tool will catalyze the use of the GoFlag408 probe set for the study of sex specific evolutionary processes or facilitate the development of sex-specific markers across the bryophyte phylogeny. We should caution that the successful use of the pipeline is contingent upon the quality of the input genome assembly and the accuracy of the markers used to identify the U and V chromosomes. As more and better assembled genomes become available, especially males, we anticipate that additional sex-specific markers will be verified, and the tool will be useful for additional lineages of mosses, as well as liverworts and hornworts.

## AUTHOR CONTRIBUTION

SK and SFM conceived of the project. SK carried out the data curation, and JLB provided additional resources. SK designed and implemented the pipeline, with input from ELB. SK carried out the formal analyses and prepared the visualizations. SK and SFM wrote the original draft of the manuscript, and all authors contributed to the review and editing of the final manuscript.

## ACKNOWLEDGEMENT

We thank the McDaniel lab, in particular Niko Matanov, for their feedback throughout the development of this pipeline. Work in S.F.M.’s laboratory was funded by NASA Biological and Physical Sciences Division, Grant Number: SC 3786; NSF Division of Arctic Sciences, Grant Number: 2000649; and the University of Florida Astraeus Space Institute. J.L.B. is funded by the Australian Research Council Centre of Excellence for Plant Success in Nature and Agriculture (CE200100015).

## DATA AVAILABILITY STATEMENT

UVfinder is available on GitHub: https://github.com/PF03106/UVfinder. All inferred gene trees and species trees are deposited in the GitHub.

## TITLES FOR SUPPORTING INFORMATION

Figure S1. Gene tree of *G5892*. *Orthothecium rufescens* and *Aulacomnium turgidum* have high topological conflict and they are highlighted in green. Branch support value in red, under each node.

Figure S2. Chromosomal distribution of target loci and identification of sex chromosomes across *Sphagnun* species. Karyotype plots showing the physical distribution of GoFlag408 target loci mapped across nine *Sphagnum* genomes. Light gray bars represent genomic scaffolds or chromosomes scaled in megabases (Mbp). Blue vertical bars indicate GoFlag loci that are not sex-linked. Regions outlined in solid pink boxes denote sex chromosomes (or sex-linked scaffolds) newly identified by the Uvfinder pipeline, while dashed pink boxes indicate sex chromosomes identified in previous studies. The bright pink marker highlights the specific GoFlag locus (*G4603*), which exhibits sex-linkage consistently in *S. balticum* and *S. palustre*. Asterisks (*) represent sex chromosome unidentified by UVfinder and previous studies.

Figure S3. Genomic and phylogenetic evidence of recent polyploidy in *Isopterygiella pulchella*. Top panels: Chromosomal distribution of sex-linked markers across scaffolds of Pseudotaxiphyllum elegans (n=10+U) and Isopterygiella pulchella (n=22). Gray horizontal bars represent individual scaffolds or chromosomes scaled in megabases (Mbp). Gery colored bars denote GoFlag query set hits, with colors indicating the number of sharing samples in Hypnales (1 to 11). Outlines signify BLAST hits to key sex-markers (female sex marker 1st-best hits in solid pink, 2nd-best hits in dashed pink, and male sex marker hits in dotted blue). The Circos plot (top right) illustrates inter-chromosomal synteny and genomic location of the duplicated sex-linked locus G5892 (paralogs A1, A2, and A3, A meaning an autosome) across the I. pulchella genome. Bottom panels: Maximum likelihood gene trees examples (G5892, G5634, and G4976) demonstrating the evolutionary timing of duplication. Blue shaded boxes highlight clusters where I. pulchella exhibits paralogs (e.g., A1, A2, A3, or U/A). Numbers below branches indicate bootstrap support values, and scale bars represent substitutions per site.

Table S1. Example of moss config/samples.tsv.

Table S2. Example of liverworts config/samples.tsv and Uvfinder assigned sex chromosome and the number of sex-linked genes.

Table S3. Example of hornworts config/samples.tsv and Uvfinder assigned sex chromosome and the number of sex-linked genes.

Table S4. Chromosome size and the number of genes on each chromosome in each of the hornwort samples.

