## Supplementary figures and images for "UVfinder: a tool to extract bryophyte sex-linked gene copies from the GoFlag408 probe set"

### Figure_S1

Supplementary Figure 1

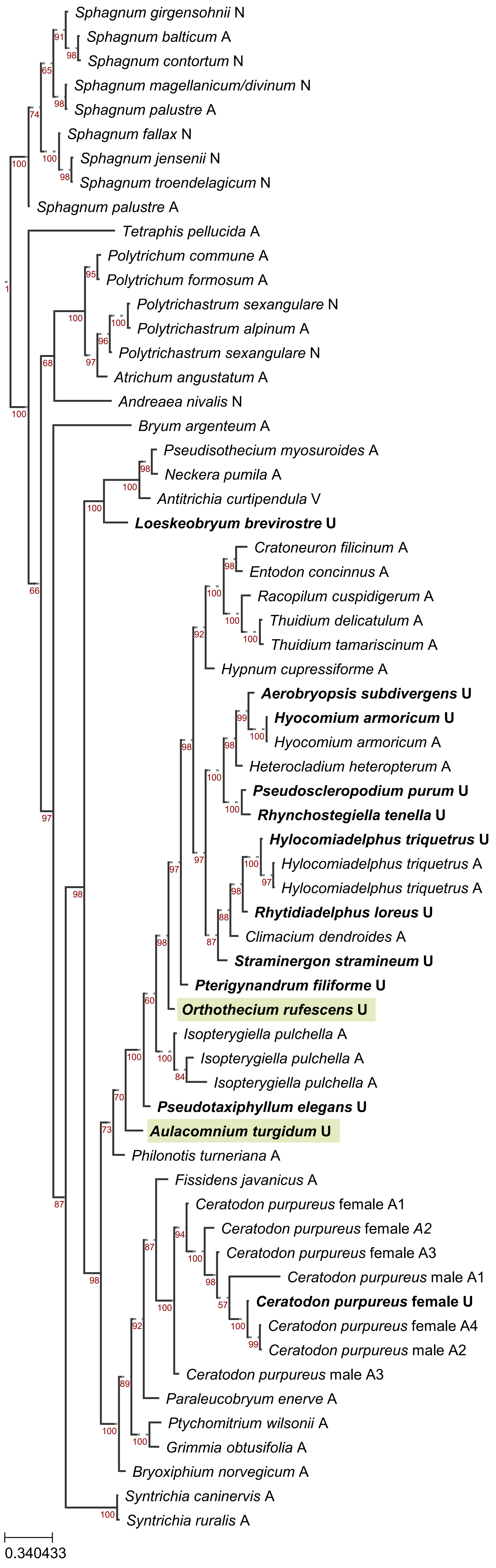

### Figure_S2

##### Supplementary Figure 2

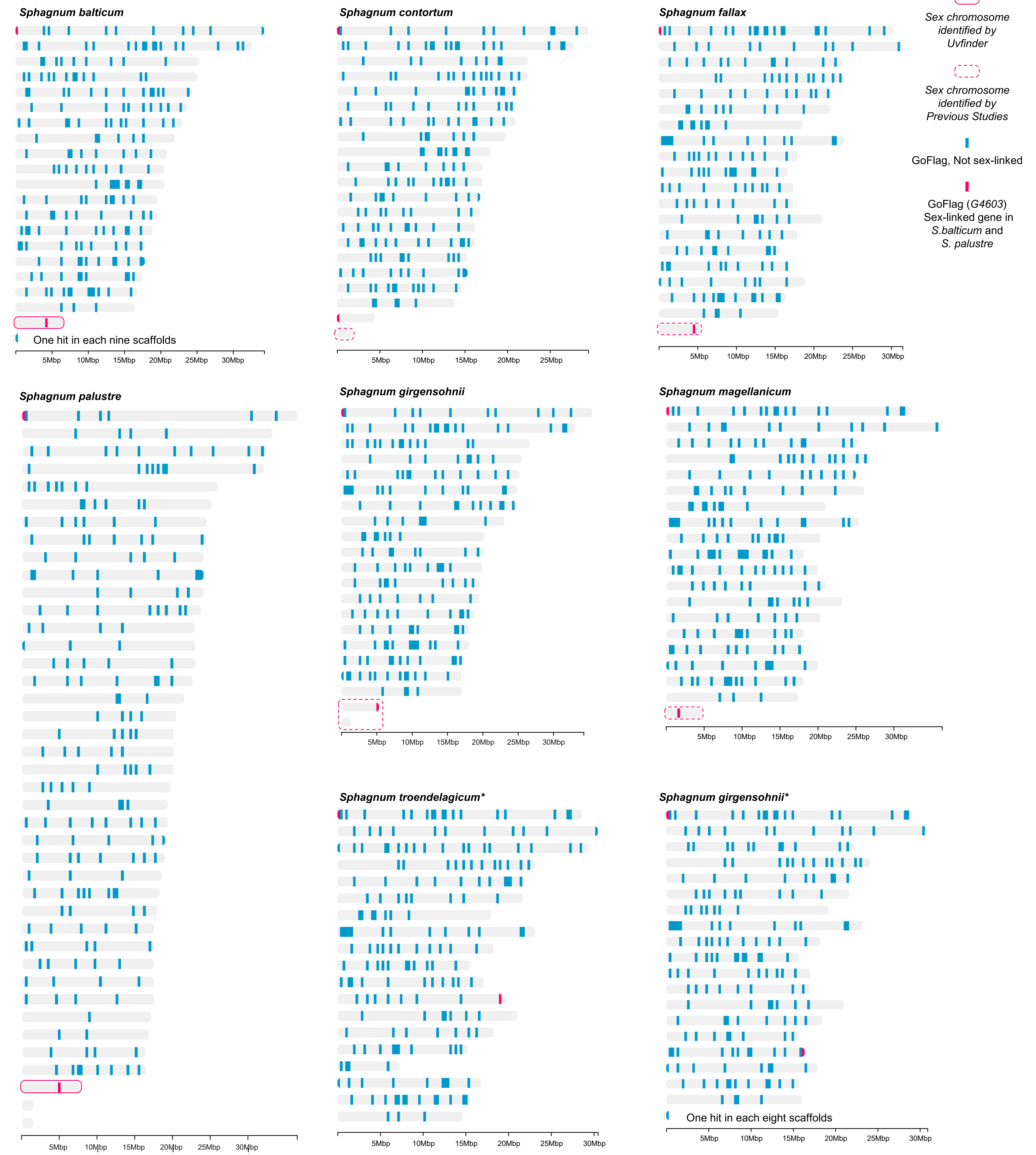

### Figure_S3

#### Supplementary Figure 3

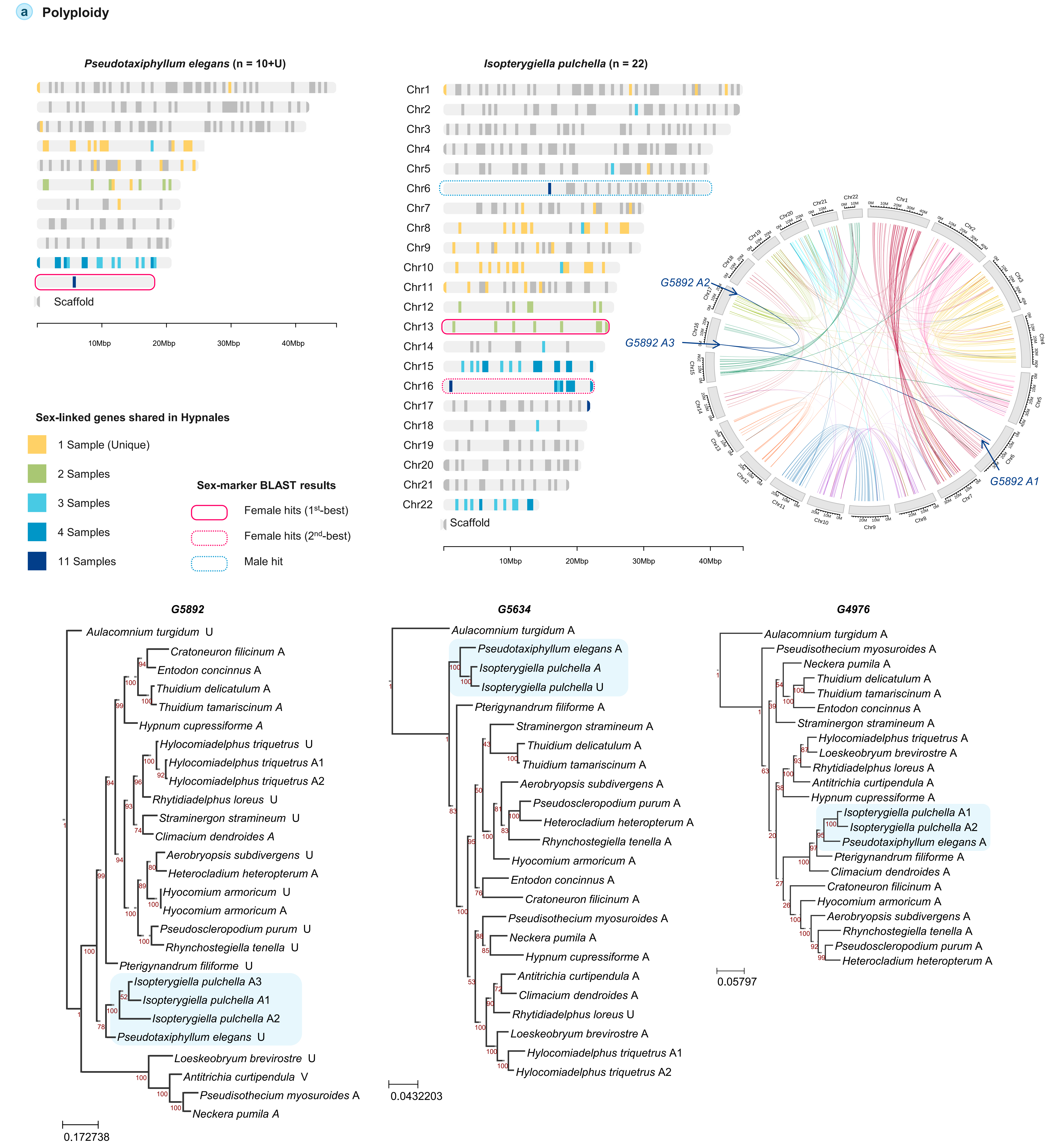
